# Atrial natriuretic peptide orchestrates a coordinated physiological response to fuel non shivering thermogenesis

**DOI:** 10.1101/866277

**Authors:** Deborah Carper, Marine Coue, Emmani Nascimento, Valentin Barquissau, Damien Lagarde, Carine Pestourie, Claire Laurens, Justine Vily Petit, Maud Soty, Laurent Monbrun, Marie-Adeline Marques, Yannick Jeanson, Yannis Sainte-Marie, Aline Mairal, Sébastien Dejean, Geneviève Tavernier, Nathalie Viguerie, Virginie Bourlier, Frank Lezoualc’h, Audrey Carrière, Wim H.M. Saris, Arne Astrup, Louis Casteilla, Gilles Mithieux, Wouter van Marken Lichtenbelt, Dominique Langin, Patrick Schrauwen, Cedric Moro

## Abstract

Atrial natriuretic peptide (ANP) is a cardiac hormone controlling blood volume and arterial pressure in mammals. It is unclear whether and how ANP controls cold-induced thermogenesis *in vivo*. Here we show that acute cold exposure induces cardiac ANP secretion in mice and humans. Genetic inactivation of ANP promotes cold intolerance and suppresses about half of cold-induced brown adipose tissue (BAT) activation in mice. While white adipocytes are resistant to ANP-mediated lipolysis at thermoneutral temperature in mice, cold exposure renders white adipocytes fully responsive to ANP to activate lipolysis and a thermogenic program, a physiological response which is dramatically suppressed in ANP null mice. ANP deficiency also blunts liver triglycerides and glycogen metabolism thus impairing fuel availability for BAT thermogenesis. ANP directly increases mitochondrial uncoupling and thermogenic genes expression in human white and brown adipocytes. Together, these results indicate that ANP is a major physiological trigger of BAT thermogenesis upon cold exposure in mammals.

## Introduction

Warm blooded animals have acquired the ability to maintain their core body temperature constant in fluctuating temperature environments through an adaptive physiological process called thermogenesis. Brown adipose tissue (BAT) is considered the major site of non-shivering thermogenesis and heat production, which allows rodents to maintain euthermia at temperatures below thermoneutrality ^1^.

Thermogenic fat cells include brown and beige adipocytes, cells that play a critical role in defending against hypothermia, obesity, and diabetes through dissipating chemical energy as heat in part through mitochondrial uncoupling protein 1 (UCP1) ^2^. Thermogenic genes can be readily induced in brown and beige adipocytes within white adipose tissue (WAT) in response to cold exposure ^3–5^. When activated, thermogenic adipocytes consume large amount of circulating triglycerides, glucose and non esterified fatty acid (NEFA) ^6^. The long-standing prevailing view is that cold sensation is primarily transmitted through the sympathetic nervous system. BAT and WAT are innervated by sympathetic fibers ^7^, which, upon cold exposure, release norepinephrine to acutely activate thermogenesis and lipolysis. Norepinephrine activates β_3_-adrenergic receptors and cAMP-dependent protein kinase (PKA) that elicit a signaling cascade via p38 mitogen-activated protein kinase (p38 MAPK) and peroxisome proliferator-activated receptor (PPAR)-γ co-activator 1α (PGC1α) to increase the transcription of UCP1 and thermogenic genes in brown adipocytes ^8^. In WAT, β_3_-adrenergic signaling activates adipocyte lipolysis through adipose triglyceride lipase (ATGL) and hormone-sensitive lipase (HSL). WAT lipolysis is essential to provide fatty acid (FA) fuels in the fasting state to sustain high respiration rates in BAT ^9, 10^.

Yet, recent studies indicate that β_3_-adrenergic receptor is dispensable in brown and beige adipocyte for cold-induced transcriptional activation of thermogenic genes in mice ^11^. Studies in mice lacking all three β-adrenergic receptors (so-called β-less mice) inferred the existence of non-adrenergic signaling pathways contributing to brown/beige adipocyte recruitment and activation ^7, 12^. Studies over the past few years identified a variety of circulating factors and hormones that may control beige adipocyte development of which cardiac natriuretic peptides (NP) are of particular interest. NP control blood volume and pressure in mammals ^13^, and were shown to be potent lipolytic hormones in human WAT (hWAT) ^14, 15^. Moreover, NP induce the transcription of *PGC1α* and *UCP1* via cGMP-dependent protein kinase (PKG) *in vitro* in human adipocytes ^16^. Chronic B-type NP (BNP) infusion in mice induces *Pgc1α* and *Ucp1* in BAT and inguinal WAT (iWAT) ^16^. However although NP induce a transcriptional thermogenic program in adipocytes, it is unclear if they are necessary and required for physiological activation of brown/beige adipocytes during acute cold exposure. Here, we show that cardiac atrial NP (ANP) is a physiological non-adrenergic activator of BAT thermogenesis through binding to its receptor guanylyl cyclase-A (GCA). We demonstrate that ANP is required for cold-induced activation of BAT thermogenesis and WAT lipolysis in mice. ANP deficiency also impairs liver triglycerides (TG) and glycogen metabolism thus contributing to reduced substrate availability to fuel BAT thermogenesis. We further show that ANP induces mitochondrial uncoupling and a thermogenic program in human primary brown adipocytes, while expression of its receptor GCA relates to a wide range of brown/beige markers and genes involved in oxidative metabolism in human subcutaneous abdominal fat. Thus, our findings identify ANP as a critical physiological endocrine regulator of non-shivering thermogenesis in mammals.

## Results

### ANP is required for non-shivering thermogenesis during acute cold exposure

[^18^F]fluorodeoxyglucose positron emission tomography-computed tomography (^18^F-FDG PET/CT) imaging indicate that acute cold exposure at 4°C for 5h induces the recruitment of several BAT depots in the neck (anterior cervical) and clavicular areas (clavicular, axillary, supraspinal, interscapular, infrascapular, anterior subcutaneous, ventral spinal and perirenal) in wild-type mice (**Fig. 1a** and **Supplementary Fig. 1a**). This is consistent with a recent study reporting similar recruitment of BAT depots in response to 7 days treatment with a β_3_-adrenergic agonist ^17^. BAT activation upon cold exposure was associated with a 3-4 fold increase of cardiac *Nppa* (ANP) expression (**Fig. 1b**) and 2-fold increase of plasma ANP (**Fig. 1c**), while no change of cardiac *Nppb* (BNP) expression (**Fig. 1d**) and plasma BNP (**Fig. 1e**) were observed. Thus, this demonstrates that ANP rather than BNP behaves as a physiological endocrine ligand of GCA in response to cold stress. Cold exposure increases blood pressure and cardiac filling pressure, which are the main physiological stimuli of cardiac ANP secretion ^18^.

**Fig. 1.**
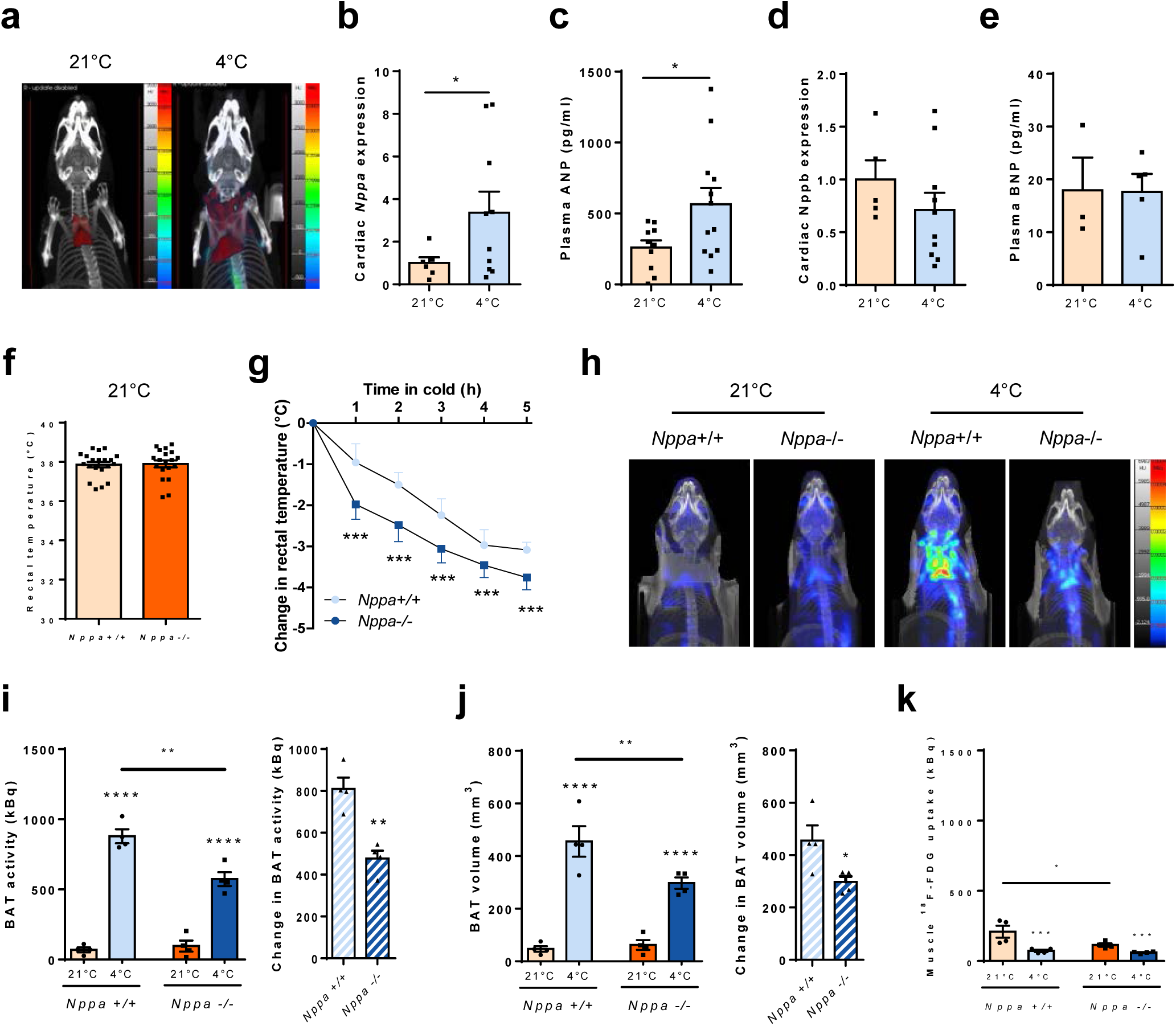
ANP is required for non-shivering thermogenesis during acute cold exposure. (a) Representative ^18^F-FDG PET/CT images of BAT recruitment around the neck in WT mice at room temperature (21°C) and exposed for 5h to 4°C (b) Relative cardiac *Nppa* (ANP) expression in WT mice housed at RT and after acute cold exposure (5h at 4°C) (n=6-10) (c) Plasma ANP levels in WT mice housed at RT and after acute cold exposure (n=10-12) (d) Relative cardiac *Nppb (BN*P) expression in WT mice housed at RT and after acute cold exposure (n=5-10) (e) Plasma *BNP* levels in WT mice housed at RT and after acute cold exposure (n=3-5) (f) Rectal temperature in *Nppa*+/+ and *Nppa*-/- mice housed at RT (n=19-20) (g) Change in rectal temperature from baseline in *Nppa*+/+ and *Nppa*-/- mice during 5h cold exposure (n=5-7) (h) Representative ^18^F-FDG PET/CT images of the neck/shoulder area indicating BAT activity in *Nppa*+/+ and *Nppa*-/- mice at RT and during cold exposure (i) Quantitative scatter plot graph and cold-induced BAT activity of *Nppa*+/+ and *Nppa*-/- mice at RT and during cold exposure (n=4) (j) Quantitative scatter plot graph and cold-induced BAT volume of *Nppa*+/+ and *Nppa*-/- mice at RT and during cold exposure (n=4) (k) Muscle ^18^F-FDG uptake of *Nppa*+/+ and *Nppa*-/- mice at RT and during cold exposure (n=4) Results are shown as mean ± SEM. *p<0.05, **p<0.01, ***p<0.001

To delineate the role of ANP in non-shivering thermogenesis, we next challenged ANP null mice (*Nppa*-/-) at 4°C. *Nppa*-/- mice had no detectable cardiac *Nppa* expression (**Supplementary Fig. 1b**) and no functional compensation by neither cardiac *Nppb* expression (**Supplementary Fig. 1c**) nor plasma BNP (**Supplementary Fig. 1d**) upon cold exposure. ANP deficiency was associated with the development of cardiac hypertrophy as shown by the increased heart weight to body weight ratio (**Supplementary Fig. 1e**) and left ventricular mass (**Supplementary Fig. 1f**). Since an intact cardiac function is required for functional non-shivering thermogenesis upon acute cold exposure ^9^, we show here that cardiac hypertrophy in *Nppa*-/- mice does not alter systolic function since left ventricular ejection fraction remains comparable to wild-type littermates (*Nppa+/+*) (**Supplementary Fig. 1g**). *Nppa*-/- mice had normal rectal temperature at room temperature (RT, 21°C) (**Fig. 1f**) and at thermoneutrality (30°C) (**Supplementary Fig. 1h**) compared with their wild-type littermates but became cold intolerant upon acute cold exposure (**Fig. 1g**). Since BAT is the main site of non-shivering thermogenesis, we measured BAT activity and recruitment by ^18^F-FDG PET/CT in *Nppa*+/+ and *Nppa*-/- mice at RT and 4°C (**Fig. 1h**). Acute cold remarkably increased BAT activity and volume, while cold-induced BAT activity (5.9 vs. 12.7 fold, p<0.01) (**Fig. 1i**) and volume (**Fig. 1j**) (4.8 vs. 9.7 fold, p<0.05) were both severely blunted in *Nppa*-/- vs. *Nppa*+/+ mice. This effect may be ascribed to the lack of ANP since expression level of genes involved in ANP signaling such as *Gca*, NP clearance receptor (*Nprc*) and cGMP-dependent protein kinase (*Prkg1*) was comparable in BAT of *Nppa*-/- and *Nppa*+/+ mice (**Supplementary Fig. 1i**). Fractional ^18^F-FDG uptake by the hind limb muscle quadriceps was reduced in *Nppa*-/- mice at 21°C, and very low at 4°C (<1% of BAT) but remained unchanged between both genotypes (**Fig. 1k**). Since β-adernergic receptors ^12^ (**Supplementary Fig. 1j, k, l**) and adenosin receptors ^19^ (**Supplementary Fig. 1m, n**) have been linked to BAT thermogenesis, we measured their gene expression levels and did not find significant changes between genotypes. Altogether, ANP deficiency severely blunts physiological BAT activation and recruitment.

### ANP-deficiency induces BAT morphological and molecular changes

Histological analysis of BAT morphology revealed the presence and accumulation of multiple and large lipid droplets in *Nppa*-/- compared with *Nppa*+/+ mice after cold exposure (**Fig. 2a**). The BAT morphology of *Nppa*-/- mice at 4°C resembles the one observed at thermoneutral temperature (30°C) in both genotypes (**Supplementary Fig. 2a**) which is characteristic of a dysfunctional BAT. Since BAT is a highly vascularized tissue ^2^ and GCA activation can modulate angiogenesis in some vascular beds ^20^, we next investigated BAT vascularization through lectin staining. No visual difference in BAT capillary density was observed between *Nppa*-/- and *Nppa*+/+ (**Supplementary Fig. 2b**). Compared to what is observed after 4°C exposure, no visual difference in BAT morphology (**Supplementary Fig. 2a**) and expression levels of thermogenic genes (**Supplementary Fig. 2c**) were noted at RT between *Nppa*-/- and *Nppa*+/+ mice. In contrast, cold-induced *Ucp1* (**Fig. 2b**) and *Pgc1α* (**Fig. 2c**) gene expression was severely blunted in *Nppa*-/- mice. We further confirmed a significant reduction of UCP1 protein content in interscapular BAT (iBAT) at 4°C in *Nppa*-/- mice (**Fig. 2d**). Consistent with an impaired cold-induced BAT activation, we observed a blunted cold-mediated response of two canonical PPAR-target genes ^21, 22^, *e.g.* carnitine palmitoyl transferase-1B (*Cpt1b*) (**Fig. 2e**) and perilipin 2 (*Plin2*) (**Fig. 2f**) in *Nppa*-/- mice. Previous work has shown that PGC1α, once induced by acute cold, co-activates PPARγ, a crucial nuclear receptor orchestrating the transcriptional program for substrate oxidation and thermogenesis in BAT ^23^. Brown fat lipid accumulation in *Nppa*-/- mice seems to occur independently of changes in NEFA transport through *Cd36* (**Fig. 2g**) and lipoprotein lipase (*Lpl*) (**Fig. 2h**) gene expression which were similarly induced by cold exposure in both genotypes. In the same line, no change in protein content of the rate-limiting enzymes ATGL (**Fig. 2i**) and HSL (**Fig. 2j**), as well as *de novo* lipogenesis genes such as carbohydrate responsive-element binding protein-β (*Chrebpβ*), acetyl-coA carboxylase 1 (*Acc1*) and fatty acid synthase (*Fas*) (**Supplementary Fig. 2d, e, f**) were noted in *Nppa*-/- versus wild-type control. Reduced cold-induced BAT activation and glucose uptake were associated with a significant increase of cold-induced *Glut1* expression (**Supplementary Fig. 2g**) while no change in *Glut4* (**Supplementary Fig. 2h**) were observed in *Nppa*-/- mice.

**Fig. 2.**
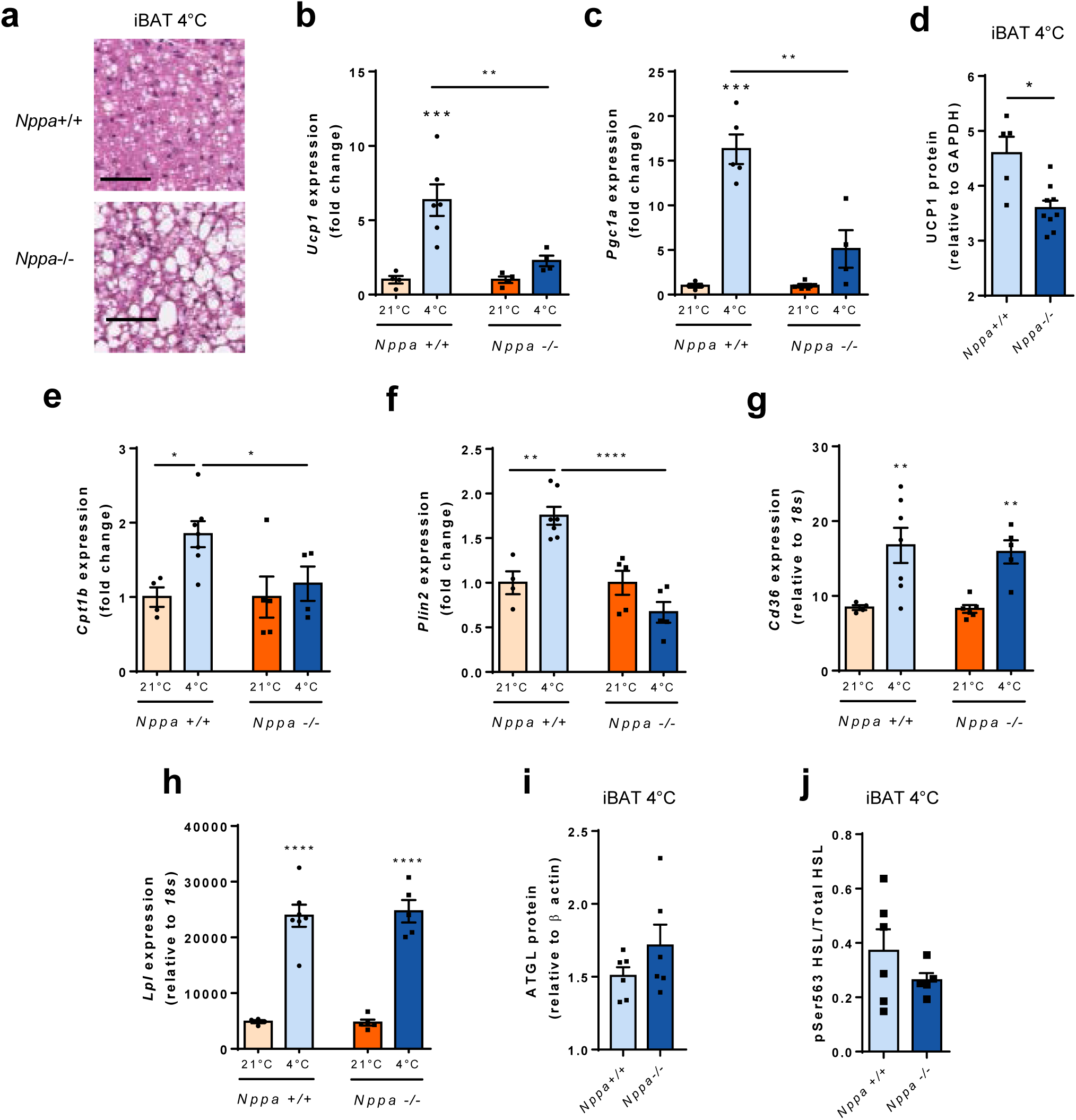
ANP-deficiency induces BAT morphological and molecular changes. (a) Representative Hematoxylin/Eosin staining of interscapular BAT (iBAT) sections of *Nppa*+/+ and *Nppa*-/- mice after cold exposure. Scale bar = 50µm (b-c) Relative mRNA levels of *Ucp1* (b) and *Pgc1α* (c) in iBAT from *Nppa*+/+ and *Nppa*-/- mice housed at RT or after cold exposure (n=4-6) (d) Relative UCP1 protein content in iBAT of *Nppa*+/+ and *Nppa*-/- mice after cold exposure (n=5-9) (e-f-g-h) Relative mRNA levels of *Cpt1b* (e), *Plin2* (f), *Cd36* (g) and *Lpl* (h) in iBAT from *Nppa*+/+ and *Nppa*-/- mice housed at RT or after cold exposure (n=4-7) (i) Relative ATGL protein content in iBAT of *Nppa*+/+ and *Nppa*-/- mice after cold exposure (n=6) (j) Relative pS563 HSL protein content in iBAT of *Nppa*+/+ and *Nppa*-/- mice after cold exposure (n=5-6) Results are shown as mean ± SEM. *p<0.05, **p<0.01, ***p<0.001, ****p<0,0001

Collectively, ANP deficiency causes marked morphological and cellular alterations of BAT biology and impairs cold-induced thermogenic genes activation which seems independent of changes in FA uptake and TG hydrolysis.

### ANP is required for beige adipocyte recruitment during acute cold exposure

Previous studies have shown that the ratio of GCA-to-NPRC expression determines cardiac NP biological activity in human and mice adipose tissue (AT) ^24–27^. Thus, genetic ablation of adipose NPRC increases NP signaling through GCA in mice ^22^. Here, we show that acute cold exposure induces WAT changes in NP receptor expression compared to mice housed at thermoneutral temperature, in a depot-specific manner. Acute cold exposure up-regulated the ratio of GCA-to-NPRC mRNA (**Supplementary Fig. 3a and Supplementary Fig. 3b**) and protein expression (**Fig. 3a**) in the three WAT depots tested. As previously described ^16^, we noted that NPRC protein was significantly diminished in iWAT (**Fig. 3b** and **Supplementary Fig. 3c**), while GCA protein remained unchanged (**Fig. 3b** and **Supplementary Fig. 3d**). Conversely, acute cold exposure up-regulated GCA protein expression specifically in epididymal WAT (eWAT) (**Fig. 3c** and **Supplementary Fig. 3d**) and retroperitoneal WAT (rpWAT) (**Fig. 3d**), while NPRC remained unaffected (**Figs. 3c, 3d** and **Supplementary Fig. 3c**). Of interest, increased *Gca* gene expression was also observed in primary mouse adipocytes exposed to cold in culture (31°C) (**Supplementary Fig. 3e**) while *Nprc* remained unchanged (**Supplementary Fig. 3f**), demonstrating a cell-autonomous increase of the GCA-to-NPRC ratio in cold-exposed white adipocytes (**Supplementary Fig. 3g**). Cold-induced up-regulation of GCA-to-NPRC ratio coincided with a sharp increase of p38 MAPK phosphorylation (**Fig. 3d**), and robust induction of its downstream transcriptional targets UCP1 and PGC1α in iWAT (**Supplementary Fig. 3h**), eWAT (**Supplementary Fig. 3i**), rpWAT (**Supplementary Fig. 3j**), and iBAT (**Supplementary Fig. 3k**). The induction of beige adipocytes in WAT, the so-called browning/beige-ing process, is highly adipose depot dependent in mice ^28, 29^. The iWAT and rpWAT undergo the most profound induction of UCP1 (>30 fold) whereas the eWAT exhibit a weak response (<10 fold). Cold-induced transcriptional activation of thermogenic genes was suppressed by ∼40% in iWAT (**Fig. 3e**) and eWAT (**Fig. 3f**), and severely impaired in rpWAT (**Fig. 3g**) of ANP null mice compared to wild type mice. The GCA-to-NPRC mRNA ratio was robustly induced in all WAT depots with a more pronounced induction in rpWAT (∼24 fold) compared to iWAT (∼1.75 fold) and eWAT (∼2.25 fold) upon cold exposure (**Supplementary Fig. 3b**). Moreover, rpWAT is sensitive to browning ^30^ and anatomically close to the kidneys, one main physiological site of action of ANP ^13^. Thus, our data show that ANP-mediated browning is WAT depot dependent with rpWAT being the most responsive depot. Along with PGC1α and UCP1, we observed a significant reduction of PR domain containing 16 (PRDM16) mRNA levels. The thermogenic activity of brown and beige adipocytes is conferred by a core gene program controlled by the master transcriptional regulator PRDM16 shown to physically interact with PGC1α to transactivate UCP1 transcription ^31^. Consistent with cold-induced thermogenic gene expression, wild-type mice exposed to acute cold showed augmented iWAT browning compared to mice housed at thermoneutrality as evidenced by increased emergence of smaller adipocytes containing multilocular lipid droplets and Hematoxylin/Eosin (H&E) staining, a physiological response that was suppressed in ANP null mice (**Fig. 3h**). Similarly to BAT, we did not find significant changes in the level of expression of cold-regulated genes involved in the control of beige adipocyte thermogenesis such as *β1ar* (**Supplementary Fig. 3l**), *β2ar* (**Supplementary Fig. 3m**), *β3ar* (**Supplementary Fig. 3n**) and *Serca2b* ^10^ (**Supplementary Fig. 3o**). No major change in genes involved in NP signaling and thermogenesis was observed in iWAT (**Supplementary Figs. 4a and 4b**) and eWAT (**Supplementary Figs. 4c and 4d**) at 30°C and 21°C between genotypes. No visual histomorphological differences in iWAT, eWAT and rpWAT were noted at thermoneutrality (**Supplementary Fig. 4e**) and RT (**Supplementary Fig. 4f**) between *Nppa*+/+ and *Nppa*-/- mice. Overall, our data emphasizes that upon a physiological cold stress, cardiac ANP released significantly contributes to thermogenic adipocytes activation and WAT browning.

**Fig. 3.**
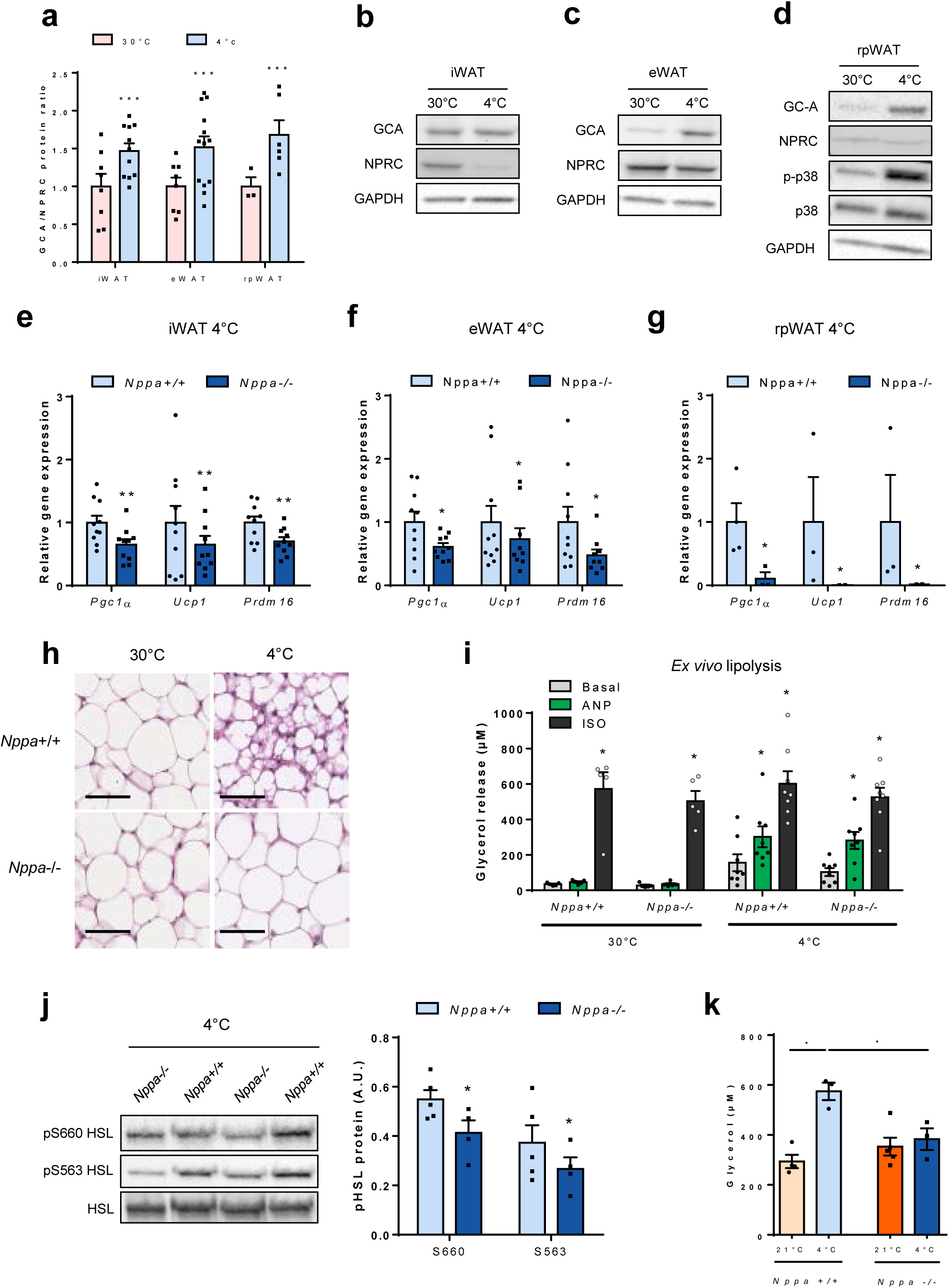
ANP is required for beige adipocyte recruitment and lipolysis during acute cold exposure. (a) GCA/NPRC protein ratio in iWAT, eWAT and rpWAT of WT mice housed at thermoneutral temperature (30°C) or after acute cold exposure (5h at 4°C) (n=6-13) (b-c) Representative immunoblot of GC-A, NPRC and GAPDH in iWAT (b) and eWAT (c) of WT mice housed at 30°C or after acute cold exposure (d) Representative immunoblot of GC-A, NPRC, p-p38, p38 and GAPDH in rpWAT of WT mice housed at 30°C or after acute cold exposure (e-f-g) Relative mRNA levels of *Pgc1a*, *Ucp1* and *Prdm16* in iWAT (n=10) (e), eWAT (n=10) (f) and rpWAT (n=3-5) (g) from *Nppa*+/+ and *Nppa*-/- mice after acute cold exposure (h) Representative Hematoxylin/Eosin staining of iWAT from *Nppa*+/+ and *Nppa*-/- mice housed at 30°C or after cold exposure. Scale bar = 50µm (i) *Ex vivo* adipocyte lipolysis in eWAT of *Nppa*+/+ and *Nppa*-/- mice housed at 30°C or after acute cold exposure under basal, ANP 10 µM and isoproterenol 1µM-stimulated conditions (n=5-8) (j) Representative immunoblot and quantitative bar graph of pS660 HSL, pS563 HSL and HSL total protein in eWAT of *Nppa*+/+ and *Nppa*-/- mice after cold exposure (n=4-5) (k) Plasma glycerol levels of *Nppa*+/+ and *Nppa*-/- mice housed at 30°C and after acute cold exposure (n=3-5) Results are shown as mean ± SEM. *p<0.05, **p<0.01, ***p<0.001

### ANP is required for white adipocyte lipolysis during acute cold exposure

Since acute exposure to 4°C up-regulates the ratio of GCA-to-NPRC protein in WAT, we hypothesized that white adipocytes would become sensitive to ANP-induced lipolysis in cold conditions. Former studies indicated that murine adipocytes are resistant to the lipolytic effect of ANP ^32^, while genetic ablation of NPRC in mice restores a lipolytic effect of ANP ^16^. Here we demonstrate that acute cold-exposure renders white adipocytes fully responsive to ANP-mediated lipolysis, whereas in mice housed at thermoneutral temperature ANP shows no lipolytic effect as previously observed. Acute cold exposure increased ANP-mediated glycerol (**Fig. 3i**) and NEFA (**Supplementary Fig. 3g**) release by 2-3 fold compared to basal conditions in adipocytes isolated from eWAT, while the effect of isoproterenol remained unchanged compared to thermoneutral temperature. The cold-mediated *ex vivo* WAT lipolytic capacity was comparable between wild-type and ANP null mice (**Fig. 3i**). This is in agreement with the lack of change of GCA (**Supplementary Fig. 4h**) and NPRC (**Supplementary Fig. 4i**) protein expression in eWAT of ANP null versus wild-type mice, indicating no functional compensation. However, cold-exposed ANP null mice had a significant down-regulation of HSL phosphorylation at Ser-660 and Ser-563 in eWAT (**Fig. 3j**), two main activating sites targeted by both PKA and PKG in response to catecholamines and ANP stimulation ^33^. This suggests that the lack of ANP *in vivo* associates with a lower cold-induced activation of HSL in eWAT of ANP null mice. This translated into a severely defective cold-induced lipolysis in ANP knockout mice as reflected by the changes in plasma glycerol concentrations between thermoneutral temperature and acute cold (**Fig. 3k**). Recent studies challenged the established view that intracellular lipolysis of lipid droplets inside BAT is rate-limiting for non-shivering thermogenesis. Rather, these studies showed that WAT lipolysis is essential to fuel BAT with FA for heat production during fasting ^9, 10^. Together, this suggests that the blunted cold-induced lipolytic response could contribute to the observed defective BAT activity and thermogenesis of ANP null mice. Altogether our data stress that physiological release of ANP during cold exposure induces lipolysis to fuel BAT thermogenesis.

### ANP deficiency impairs plasma triglycerides and glucose responses to cold

Previous work suggested that, besides NEFA, circulating TG and glucose are major substrates for BAT thermogenesis ^6^. In this study, we found reduced blood glucose levels upon cold exposure after 2h (**Supplementary Fig. 5a**), while observing a time-dependent reduction of circulating insulin (**Supplementary Fig. 5b**) and circulating TG levels (**Supplementary Fig. 5c**) during a time-course of cold exposure. We further observed decreased levels of circulating TG (**Fig. 4a**) and a blunted cold-induced plasma clearance of circulating TG in *Nppa*-/- mice (**Fig. 4b**). This occurred despite no significant difference in liver TG content between genotypes (**Fig. 4c, d**). Lipolysis-derived NEFA availability is a major determinant of liver TG production ^34^. In line with a recent study ^9^, hepatic genes related to lipid metabolism were strongly altered during cold exposure (**Fig. 4e, f** and **Supplementary Fig. 5d, g**). Cold suppressed *Cd36* (**Fig. 4e**) and *Pparα* (**Supplementary Fig. 5d**) mRNA levels, while briskly inducing *Ppargc1a* (**Supplementary Fig. 5e**), *Atgl* (**Supplementary Fig. 5f**) and *Fgf21* (**Supplementary Fig. 5g**). Of interest, altered lipolysis in ANP-deficient mice was associated with a compensatory increase of liver *Cd36* gene expression (**Fig. 4e**), while both liver *Cpt1a* mRNA levels (**Fig. 4f**) and plasma ketone bodies levels (**Fig. 4g**) were significantly reduced in *Nppa*-/- mice. This reveals a blunted NEFA utilization in the liver of ANP-deficient mice. In mirror of cold-induced NEFA utilization in liver, we observed a suppressed expression of *de novo* lipogenesis genes such as *Chrebp* (**Supplementary Fig. 5h**), *Acly* (**Supplementary Fig. 5i**), *Elovl3* (**Supplementary Fig. 5j**), and *Fas* (**Supplementary Fig. 5k**) that was similar in control and ANP deficient mice.

**Fig. 4.**
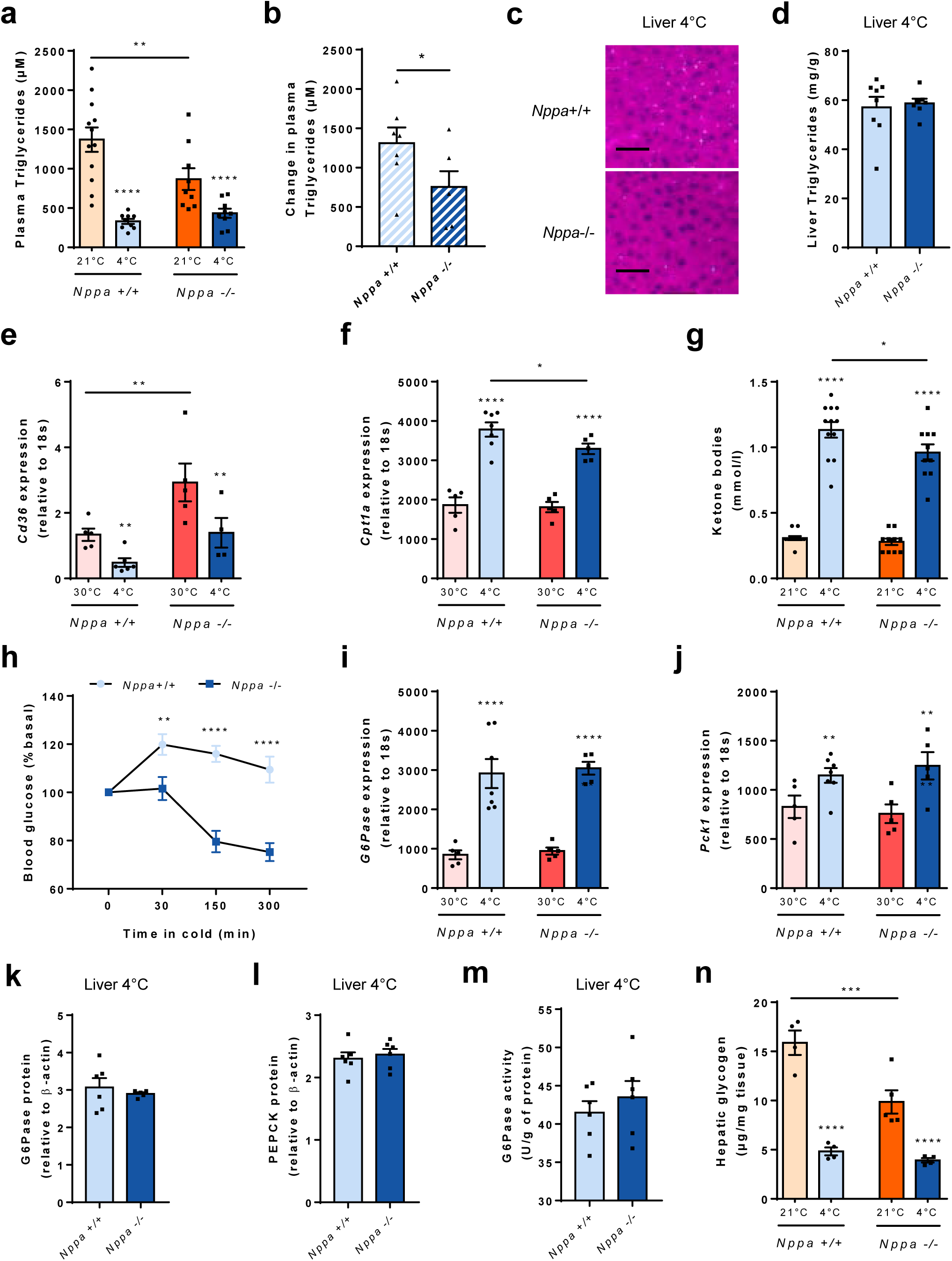
ANP deficiency impairs plasma triglycerides and glucose responses to cold. (a) Plasma triglycerides levels of *Nppa*+/+ and *Nppa*-/- mice housed at RT and after acute cold exposure (n=9-12) (b) Change in plasma triglycerides levels of *Nppa*+/+ and *Nppa*-/- mice housed at RT after acute cold exposure (n=6-7) (c) Representative Hematoxylin/Eosin staining of liver from *Nppa*+/+ and *Nppa*-/- mice after cold exposure. Scale bar = 50µm (d) Liver triglycerides content of *Nppa*+/+ and *Nppa*-/- mice after acute cold exposure (n=7-8) (e-f) Relative mRNA levels of *Cd36* (e) and *Cpt1a* (f) in iBAT from *Nppa*+/+ and *Nppa*-/- mice housed at 30°C or after cold exposure (n=4-7) (g) Plasma ketone bodies levels of *Nppa*+/+ and *Nppa*-/- mice housed at RT and after acute cold exposure (n=10-12) (h) Change in blood glucose from baseline in *Nppa*+/+ and *Nppa*-/- mice during 5h cold exposure (n=10-13) (i-j) Relative mRNA levels of *G6Pase* (i) and *Pck1* (j) in liver from *Nppa*+/+ and *Nppa*-/- mice housed at 30°C or after cold exposure (n=5-7) (k-l) Relative G6Pase (k) and PEPCK (l) protein content in liver of *Nppa*+/+ and *Nppa*-/- mice after cold exposure (n=5-6) (m) G6Pase activity in liver of *Nppa*+/+ and *Nppa*-/- mice after cold exposure (n=6) (n) Hepatic glycogen content in liver of *Nppa*+/+ and *Nppa*-/- mice housed at RT or after cold exposure (n=4-5) Results are shown as mean ± SEM. *p<0.05, **p<0.01, ***p<0.001, ****p<0,0001

Because lipolysis-derived NEFA availability is also a strong determinant of endogenous glucose production in mice ^35^, we investigated glucose metabolism in cold-exposed mice. Remarkably, ANP null mice were not able to maintain their blood glucose levels in a normal physiological range (**Fig. 4h**). This phenomenon was independent of changes in glucose-6-phosphatase (*G6pase*) (**Fig. 4i**) and phosphoenolpyruvate carboxy-kinase-1 (*Pck1*) (**Fig. 4j**) mRNA levels, protein content (**Fig. 4k, l**) and G6Pase activity (**Fig. 4m**) in liver of ANP KO mice versus control. Importantly, we observed a strong liver glycogen depletion upon acute cold exposure (∼70%) in control mice, while cold-induced glycogen depletion was strongly blunted in *Nppa*-/- mice (**Fig. 4n**), thus coinciding with the inability to maintain normal blood glucose levels during cold exposure. In summary, our data together indicate that ANP deficiency impairs liver TG and glycogen metabolism thus contributing to reduced substrate availability to fuel BAT thermogenesis.

### Cold induces ANP and GC-A is associated with brown/beige thermogenic markers in human subcutaneous abdominal WAT in humans

To further examine if ANP could play a role in cold-induced activation of BAT in humans, we determined circulating plasma NP levels in human volunteers exposed to mild cold. Thus, recent studies using ^18^F-FDG PET/CT revealed that acute cold exposure readily activates BAT in humans ^36, 37^, however the physiological cues orchestrating hBAT activation are still unclear. In a study in which 1h cold exposure at 16°C increased mean blood pressure, BAT activity and systemic lipolysis in lean healthy male volunteers (**Fig. 5a**) ^38^, we measured a significant increase of plasma ANP levels by 1.7 fold (**Fig. 5b**) whereas plasma BNP concentrations remained strictly unchanged (**Fig. 5c**). In light of the previous findings in human primary BAT-derived adipocytes, this suggests that ANP is a cold-induced endocrine activator of BAT function in humans. Thus in agreement with mice data, this implies that ANP behaves as the physiological endocrine ligand of GCA in response to cold stress in humans.

**Fig. 5.**
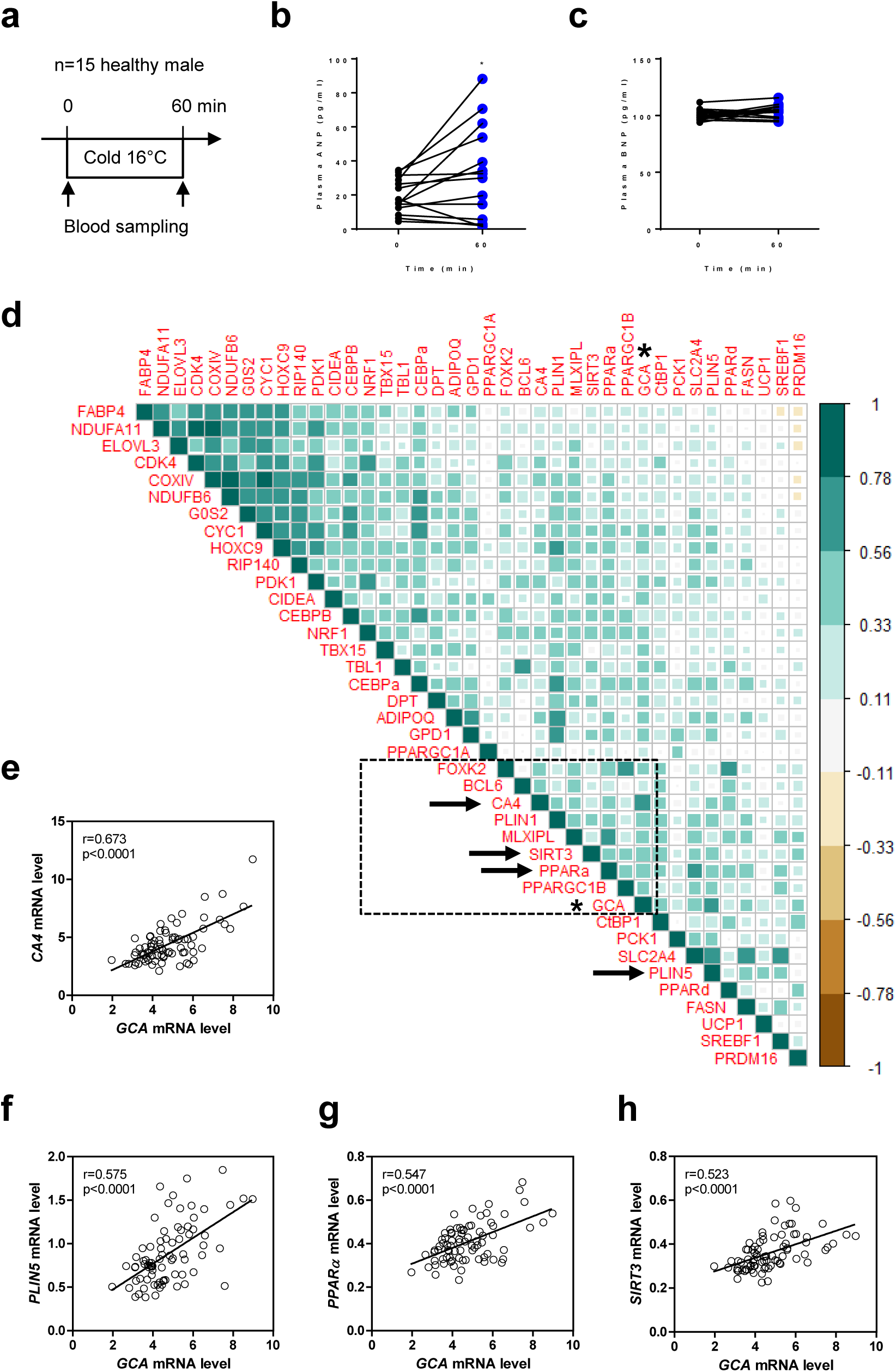
GC-A is associated with brown/beige and thermogenic markers in human subcutaneous abdominal WAT. (a) Study design of acute cold exposure in human healthy volunteers (b) Plasma ANP levels in human healthy volunteers before and after 60min cold exposure (n=14) (c) Plasma BNP levels in human healthy volunteers before and after 60min cold exposure (n=15) (d) Correlation matrix of the 39 brown/beige-specific gene markers significantly correlated with *GCA*. Color intensity and spread are directly proportional to correlation coefficients as shown by the vertical scale. The dot line box plot indicates a gene cluster containing GCA indicated by a star * (e-f-g-h) Univariate linear regression between *GCA* mRNA level and *CA4* (b), *PLIN5* (c), *PPARα* (d), and *SIRT3* (e) mRNA levels in human WAT (n=79) Results are shown as mean ± SEM. *p<0.05 vs time point 0.

There is a large variability of white fat browning/beige-ing in human individuals particularly those with obesity ^24^. We next investigated the relationship between mRNA levels of GCA and markers of browning/beige-ing in subcutaneous abdominal adipose tissue in a cohort of middle-aged individuals with a wide range of body mass index selected for a high baseline expression of UCP1 (n=79). Correlation matrix analysis revealed that GCA was highly correlated with previously reported brown/beige-specific markers involved in mitochondrial oxidative metabolism, mitochondrial biogenesis, glucose and FA metabolism ^24^ (**Fig. 5d**). An optimal re-ordering of the correlation matrix based on hierarchical clustering revealed that *GCA* clustered with several brown/beige-specific gene markers such as *PPARα*, Sirtuin 3 (*SIRT3*), Carbonic Anhydrase 4 (*CA4*), Forkhead Box K2 (*FOXK2*) and PPARγ co-activator 1β (*PPARGC1B*) (dotted line box **Fig. 5d**). The top-ranking genes displaying the highest correlations with *GCA* were the beige-specific marker *CA4* (**Fig. 5e**), the lipid droplet-associated protein Perilipin 5 (*PLIN5*) (**Fig. 5f**), the transcription factor *PPARα* (**Fig. 5g**), and the brown-specific mitochondrial *SIRT3* (**Fig. 5h**), all genes highly expressed in BAT and involved in metabolic pathways supporting thermogenic function ^24^. Thus correlations between *GCA* and brown/beige-specific markers support a role for ANP/GCA signaling in *bona fide* thermogenic adipocytes of human subcutaneous abdominal WAT.

### ANP activates mitochondrial uncoupling in BAT and a thermogenic program in human primary brown/beige and white adipocytes

^18^F-FDG PET/CT revealed the existence of active BAT in the supraclavicular and neck areas of adult humans, that can be readily activated by cold exposure ^36, 39–41^. It was suggested that these human BAT (hBAT) depots are in fact composed of UCP1-positive adipocytes bearing a transcriptional signature of beige rather than brown adipocytes as found in rodents ^5^. Here we differentiated adipocytes derived from hBAT (prevertebral) or human WAT (hWAT) (neck) *in vitro* from the same subject using four independent donors as previously described ^5, 42^. mRNA expression levels of *UCP1*, *PRDM16*, Cell Death-Inducing DFFA-Like Effector A (*CIDEA)* and Nuclear respiratory factor 1 *(NRF1)* were significantly higher in brown/beige compared to white adipocytes (**Supplementary Fig. 6a**). Thus, we also observed a higher gene expression level of the NP-signaling components *NPRC*, *PRKGI* and Phosphodiesterase 5A *(PDE5A)* in BAT biopsy-derived adipocytes (**Supplementary Fig. 6b**). We next measured mitochondrial oxygen consumption under ATP synthase inhibition by oligomycin treatment to focus on mitochondrial uncoupled respiration (state 4). We have previously shown that β-adrenergic stimulation leads to pronounced mitochondrial uncoupling in human primary brown/beige adipocytes which is markedly less in human primary white adipocytes, illustrating the unique feature of human adipocytes derived from the neck region ^42^. Interestingly, we here show that ANP dose-dependently activates uncoupled mitochondrial respiration to about 50% of the effect of norepinephrine (NE) in human brown/beige adipocytes (**Fig. 6a, b**). This effect was markedly lower in WAT-derived adipocytes (**Fig. 6c, d**). ANP at the lowest dose of 100 nM nearly doubled maximal uncoupled respiration measured under carbonilcyanide p-triflouromethoxyphenylhydrazone (FCCP) in human brown/beige adipocytes (**Fig. 6e**). A similar but weaker effect was observed in hWAT adipocytes (**Fig. 6f**). This reveals that ANP can directly activate mitochondrial uncoupling and respiration in human primary brown/beige adipocytes. We next examined if ANP could induce a transcriptional thermogenic program in human brown/beige and white primary adipocytes as observed *in vivo* in mice. ANP treatment briskly increased mRNA levels of *UCP1* both in human WAT and BAT adipocytes (**Supplementary Fig. 6c**). A peak was observed after 3h treatment with levels returning to baseline after 48h and 72h treatment for *UCP1* and *CPT1B* in human brown/beige adipocytes (Supplementary Fig. 6d, e) and white (**Supplementary Fig. 6f, g**) adipocytes. Interestingly, acute treatment (3h) with ANP at 100 nM increased to a variable degree (from 1.5 to 50-fold) a number of brown markers such as *UCP1* and deiodinase type 2 (*DIO2*), beige markers such as Cbp/P300 Interacting Transactivator With Glu/Asp Rich Carboxy-Terminal Domain 1 (*CITED1*) and Elongation Of Very Long Chain Fatty Acids Protein 3 (*ELOVL3*), and mitochondrial oxidative metabolism markers *CPT1B*, NADH:Ubiquinone Oxidoreductase Subunit B6 (*NDUFB6*) and Transcription Factor A, Mitochondrial (*TFAM*) in brown/beige adipocytes (**Fig. 6g**) and white adipocytes (**Fig. 6h**). Taken together, these results indicate that ANP has the capacity to activate a thermogenic program in human primary brown/beige and white adipocytes, and mitochondrial uncoupling in brown/beige adipocytes. This implies that ANP mimetics and/or pharmacological compounds able to increase ANP/GCA-signaling may be attractive strategies to activate BAT in humans.

**Fig. 6.**
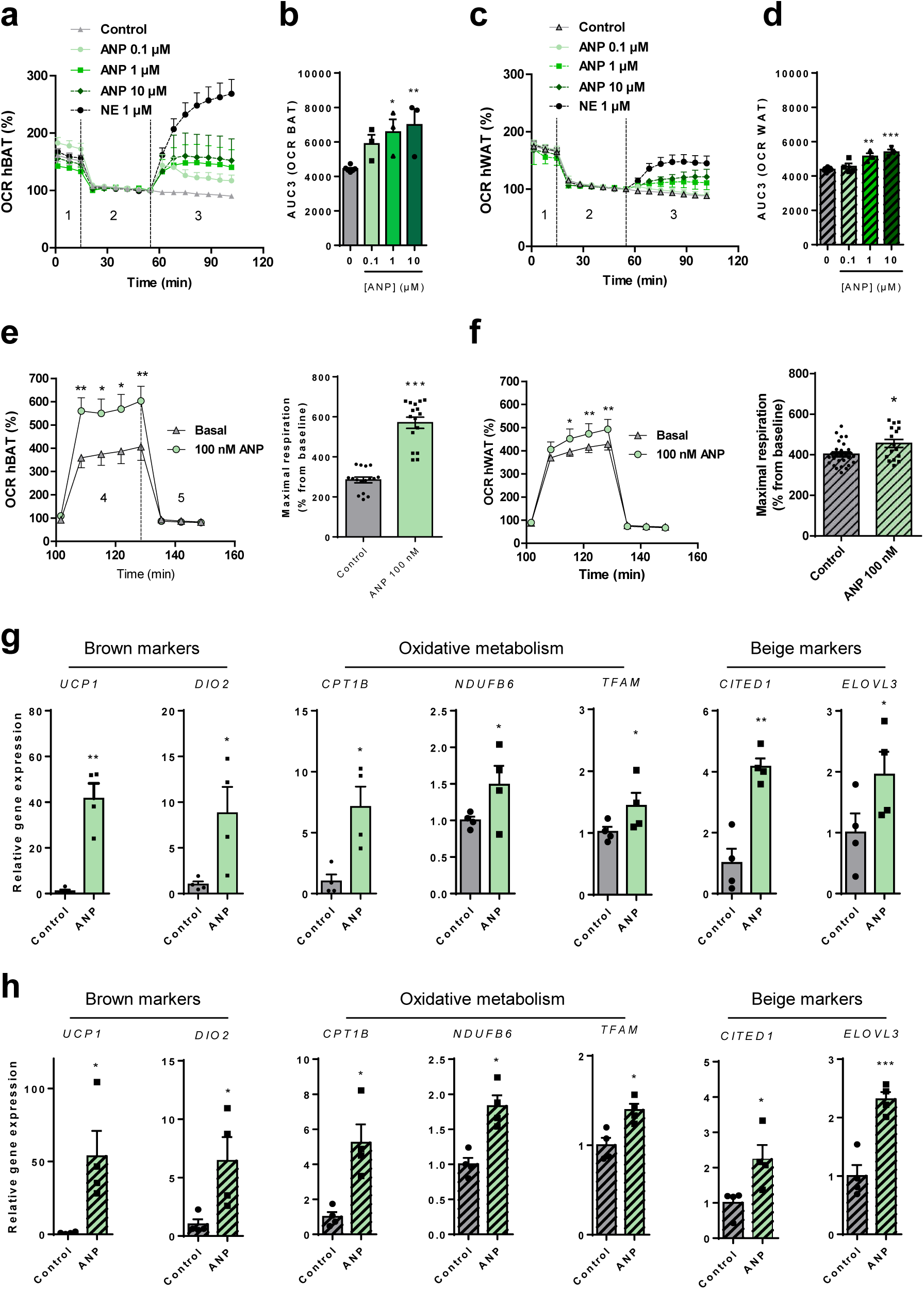
ANP activates mitochondrial uncoupling in BAT and a thermogenic program in human primary brown/beige and white adipocyte. (a) Oxygen consumption rate (OCR) of human primary brown/beige adipocytes (hBAT) in absence (control), or presence of ANP 0.1µM, ANP 1µM, ANP 10µM and NE 1µM for 3 hours (n=4). 1: basal respiration, 2: oligomycin, 3: treatments with different doses of ANP and NE (b) Area under the curve (AUC) of treatment-induced OCR calculated during phase 3 (uncoupled respiration during oligomycin inhibition of ATP synthase) in (A) (c) OCR of human primary white adipocytes WAT (hWAT) in absence (control) (n=6), or presence of ANP 0.1µM, ANP 1µM, ANP 10µM and NE 1µM for 3 hours (n=4). 1: basal respiration, 2: oligomycin, 3: treatments with different doses of ANP and NE (d) AUC of treatment-induced OCR calculated during phase 3 in (uncoupled respiration during oligomycin inhibition of ATP synthase) (C) (e) Maximal OCR (induced by FCCP) of hBAT in absence (control) or presence of ANP 100nM (n=4 independent donors). 4: FCCP, 5: rotenone and antimycin A (f) Maximal OCR (induced by FCCP) of hWAT in absence (control) or presence of ANP 100nM (n=4 independent donors). 4: FCCP, 5: rotenone and antimycin A (g) Relative mRNA levels of brown (*UCP1*, *DIO2*), brown/beige (*CPT1B*, *NDUFB6*, *TFAM*), and beige (*CITED1* and *ELOVL3*) markers in hBAT in absence (control) or presence of ANP 100nM (n=4) (h) Relative mRNA levels of brown (*UCP1*, *DIO2*), brown/beige (*CPT1B*, *NDUFB6*, *TFAM*), and beige (*CITED1* and *ELOVL3*) markers in hWAT in absence (control) or presence of ANP 100nM (n=4) Results are shown as mean ± SEM. *p<0.05, **p<0.01, ***p<0.001 vs control.

## Discussion

The current classical view in mammals is that brown/beige fat is primarily activated by norepinephrine released from sympathetic nerves upon cold exposure. However, the fact that the non selective β-agonist isoproterenol fails to activate BAT in humans ^43^ combined to the observation that β_3_-adrenergic receptor is dispensable for cold-induced thermogenic gene activation in mice ^11^, points toward the existence of alternative non-adrenergic regulatory systems that control BAT activation and function in response to cold. We here show that ANP, a cardiac hormone controlling blood volume and pressure, is necessary and required for cold-induced brown/beige adipocyte activation (Fig. 7). We further show that ANP is physiologically released upon cold exposure and activates mitochondrial uncoupling and a thermogenic program in human brown/beige adipocytes.

**Fig. 7.**
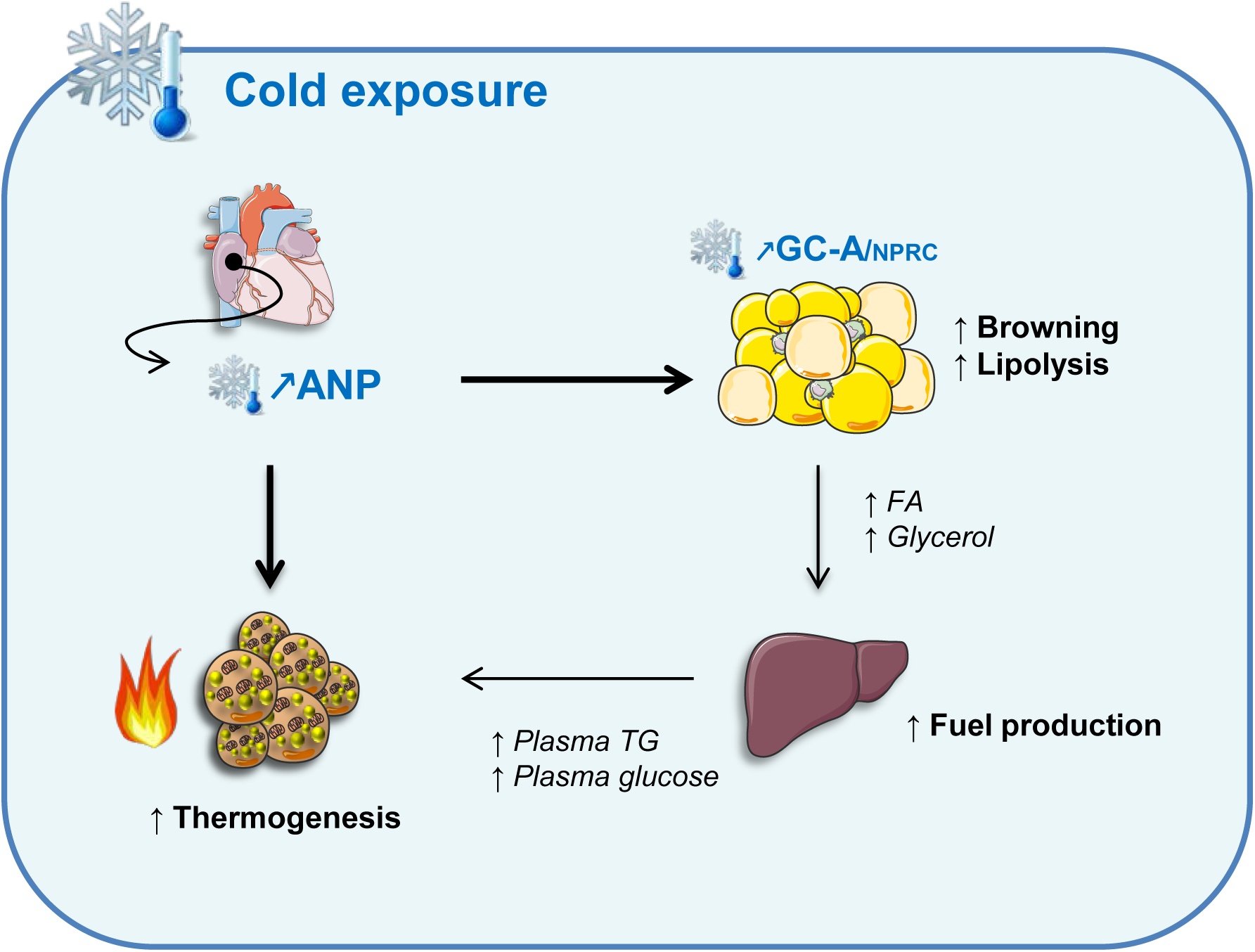
ANP triggers cold-induced brown/beige and white fat activation. Acute cold exposure increases cardiac preload and promotes ANP secretion by the heart. Cold exposure also briskly enhances ANP signaling in WAT through an increase of the GCA-to-NPRC ratio. ANP through GCA activates BAT thermogenesis and induces a transcriptional thermogenic program in WAT as well as lipolysis. FA and glycerol release by adipose tissue fuels liver TG and glucose production which are main circulating substrates of BAT to sustain heat production.

Previous work demonstrated that ANP contributes to cold-induced diuresis in healthy humans ^44^. In response to cold, contraction of superficial blood vessels will limit heat loss, and as a consequence, blood will be shunted away toward deeper large blood vessels that will increase cardiac filling pressure of the right atrium, *i.e.* increased cardiac preload. As a result, ANP secretion will be induced to normalize the increase in cardiac preload by enhancing diuresis. Herein, we demonstrate that ANP released upon cold exposure, but not BNP, will activate BAT to produce heat and maintain euthermia. BNP, the product of *Nppb*, is marginally expressed in the right atria of the heart compared to ANP ^13^. This likely explains why cardiac BNP expression and circulating levels are poorly affected by changes in cardiac filling pressure such as induced by cold exposure in this study.

In previous studies, the complete lack of all three β-adrenergic receptor (β-less mice)^12^ or sympathetic innervations in WAT through pharmacological ablation by 6-hydroxydopamine ^45^ or genetic invalidation of tyrosine kinase receptor-A ^7^ was shown to suppress partially but not completely cold-induced UCP1 expression. Although β-less mice develop hypothermia, it is still unclear to what extent BAT activity is hampered by the lack of β-adrenergic receptors. In this study, we show for the first time using ^18^F-FDG PET/CT that the lack of ANP abrogates about half of BAT activity and >60% of transcriptional activation of *Ucp1* and *Pgc1α* in BAT in response to acute cold exposure. The accumulation of multiple lipid droplets in cold-exposed BAT, *i.e.* BAT steatosis, of ANP null mice is a sign of dysfunctional BAT as observed in other mouse models ^9, 12, 31^. A failure to adequately activate BAT and UCP1 in ANP-deficient mice will lead to fat storage within lipid droplets in face of increased FA supply. This phenomenon is not observed at room temperature for which the cold stress represents a too moderate challenge to unmask prototypical BAT-related phenotypes.

Recent studies indicate that BAT activation upon cold exposure is intimately linked to WAT lipolysis in fasted mice ^9, 10^, thus highlighting the need for a thermogenic factor to activate lipolysis. Although ANP is a powerful lipolytic hormone in human adipocytes, previous studies could not reveal a lipolytic effect of ANP in mouse adipocytes ^32^. We here reconcile these studies demonstrating that cold exposure briskly increases ANP receptor GCA gene and protein expression in WAT, thus rendering mouse adipocytes responsive to ANP-mediated lipolysis. Importantly, we here demonstrate a cell-autonomous up-regulation of GCA in primary mouse WAT adipocytes cultured at 31°C instead of 37°C. This indicates that acute cold is sufficient to up-regulate GCA expression in white adipocytes independently of systemic neuro-endocrine factors. We next show that cold-induced systemic lipolysis, reflected by increased plasma glycerol levels, and HSL activation by phosphorylation in eWAT is blunted in ANP null mice.

Besides NEFA derived from WAT lipolysis, circulating TG have been shown as major BAT substrates during cold exposure ^6^. We here unravel a strong defect in plasma TG clearance in *Nppa*-/- mice during cold exposure despite a robust induction of *Lpl* in BAT of cold-exposed *Nppa*-/- mice similar to control littermates. In line with a recent study^9^, we observed drastic changes in expression of lipid metabolism genes in liver of cold-exposed mice. Cold exposure turns on FA oxidative gene networks while down-regulating lipid synthesis gene programs such as *de novo* lipogenesis. This leads to substantial FA utilization by the liver thus producing ketone bodies. Remarkably, cold-induced up-regulation of *Cpt1a*, a rate-limiting enzyme in mitochondrial FA oxidation, and ketone bodies production was impaired in *Nppa*-/- mice. This together with a reduced cold-induced lipolysis in *Nppa*-/- mice largely contributes to this observed hepatic phenotype.

Previous works also highlighted that glucose is an important substrate for BAT during cold exposure in mice ^6^ and humans ^46^. Of importance, we observed that *Nppa*-/- mice fail to maintain their blood glucose levels during acute cold exposure when compared to wild-type mice. This effect appears independent of changes in protein content and enzyme activity of PEPCK and G6Pase that are rate-limiting enzymes in hepatic glucose production. Thus this could be reasonably explained by a reduced glycogenolysis during cold exposure due to reduced liver glycogen content in *Nppa*-/- mice. In addition, a blunted ANP-mediated lipolysis in WAT during cold exposure reduces NEFA availability and therefore liver endogenous glucose production ^35^. Our findings are consistent with a previous work which showed a stimulatory effect of ANP on gluconeogenesis in perfused rat livers ^47^. Thus the absence of ANP will likely result in blunted gluconeogenesis in cold-exposed mice. Collectively, a reduced availability and utilization of circulating NEFA, TG and glucose largely contributes to impaired BAT thermogenesis in cold-exposed *Nppa*-/- mice.

In summary, we identify cardiac ANP as a physiological endocrine activator of non-shivering thermogenesis in mammals. These data uncover an intriguing evolutionary interconnection between cardiac activity and non-shivering thermogenesis. While the sympathetic nervous system remains the best-known mediator of cold-induced thermogenesis and BAT recruitment in mammals, our findings shed light on alternative pathways that have been conserved across species to maintain euthermia. They may open the path to novel pharmacological strategies targeted to enhance ANP/GCA signaling for human BAT recruitment to improve metabolic profile in individuals with obesity and type 2 diabetes.

## Methods

### Clinical studies and human subjects

#### Study 1

Ancillary study of the DiOGenes (Diet, Obesity and Genes) European Framework project (NCT00390637). For a thorough description of the overall objective and goals of this multicenter, randomized, controlled dietary intervention study, see ^48^. Briefly, the study examined the effects of dietary macronutrients on weight regain and cardiovascular risk factors. Inclusion and exclusion criteria for study participation were previously outlined ^48^. The DiOGenes study included 938 participants aged 27-63 years from 8 European countries; however, the present study used only a subgroup of 79 men and women who had high or medium *UCP1* gene expression in subcutaneous white adipose tissue as previously described ^24^ and *GCA* expression data available. Only baseline data were used in the present investigation.

#### Study 2

To investigate the effect of acute cold on circulating NP levels, each subject underwent a mild cold experiment. This experiment started with one-hour baseline measurements during thermoneutral conditions. Subsequently, subjects were exposed to one hour of mild cold exposure, in which a standardized cooling protocol was used. The mild cold experiment was conducted in a specially equipped air-permeable tent (Colorado altitude training, USA), in which ambient temperature could be tightly controlled. During baseline and the mild cold period, subjects wore standardized clothing (shorts and a t-shirt; 0.19 clo). Energy expenditure was continuously measured while body temperatures, skin perfusion (vasoconstriction) and heart rate were sampled each minute. Blood pressure was measured each 15 minutes as well as thermal comfort and thermal sensation via Visual Analog Scales (VAS). These data have been reported previously ^38^. Muscle shivering was monitored by means of EMG and VAS scales. Venous blood samples were taken during baseline and one hour after the onset of cold exposure.

### Mouse models and handling

Eight weeks old *Nppa-/-* male mice (B6.129P2-*Nppa*^tm1Unc^/J mice were backcrossed to C57BL/6J mice for at least ten generations) and their littermate control *Nppa+/+* were used. Mice were fed with a normal chow diet (Ssniff) and were housed in a pathogen-free barrier facility (12h light/dark cycle) with *ad libitum* access to water and food in standard animal care facility rooms at 21°C (RT). For cold exposure experiments, at 7 a.m. animals were placed singly and exposed for 5 hours at 4°C with water access but without food. For acclimation to thermoneutrality, mice were transferred to a chamber with controlled ambient temperature at 30°C for 4 consecutive weeks. Rectal temperature was monitored using an EcoScan Temp4/5/ thermometer (Eutech Instruments) each hour during cold exposure or at indicated time points. At the end of the protocol, mice were decapitated and blood was collected into EDTA tubes containing protease inhibitors. Organs and tissues were rapidly excised and either snap frozen in liquid nitrogen before being stored at −80°C or processed for histology. All experimental procedures were approved by our institutional animal care and use committee CEEA122 (protocol# 2016122311033178) and performed according to INSERM guidelines for the care and use of laboratory animals.

### Human primary adipocytes culture

Adipocytes derived from human BAT and WAT were obtained and differentiated as described previously ^5, 42^. The study was reviewed and approved by the ethics committee of Maastricht University Medical Center (METC 10-3-012, NL31367.068.10, NCT03111719). Informed consent was obtained before surgery. In brief, the stromal vascular fraction (SVF) was obtained from prevertebral BAT and subcutaneous WAT from the same area during thyroid surgery using a collagenase digestion. Differentiation was initiated for 7 days with differentiation medium containing biotin (33 µM), pantothenate (17 µM), insulin (100 nM), dexamethasone (100 nM), IBMX (250 µM), rosiglitazone (5 µM), T3 (2 nM), and transferrin (10 µg/ml). Cells were transferred to maintenance medium consisting of biotin (33 µM), pantothenate (17 µM), insulin (100 nM), dexamethasone (10 nM), T3 (2 nM), and transferrin (10 µg/ml) for another 5 days.

### Mouse primary adipocytes culture

SVF from iWAT was obtained from 6-week-old WT mice as previously described ^49^. iWAT was dissected, mechanically dissociated and digested for 30min at 37°C with collagenase (collagenase NB 4 Standart Grade from Coger, concentration of 0.4 U/ml diluted in proliferative medium (αMEM plus 0.25 U/ml amphotericin, 100 U/ml penicillin, 100 mg/ml streptomycin, 0.016 mM biotin, 100 μM ascorbic acid, 0.018 mM pantothenic acid and 10% new-born calf serum)). After filtration, red blood cells lysis and centrifugation, the pellet was resuspended in proliferative medium. SVF cells were then counted, plated at 10000 cells/cm² and rinsed in PBS 3 hours after plating. Cells were maintained at 37°C (5% CO_2_) and re-fed every 48h. Adherent cells were grown to 80% confluency in proliferative medium. Cells were then exposed to an adipogenic cocktail (proliferative medium supplemented with 5 μg/ml insulin, 2 ng/ml T3, 33.3 nM dexamethazone, 10 μg/ml transferrin and 1 μM rosiglitazone) and used after 8 days of differentiation.

### Cellular cooling

Fully differentiated inguinal mouse primary adipocytes were kept in a 37°C incubator with 5% CO_2_ before experiments. Before cooling treatment, medium was refreshed with prewarmed adipogenic cocktail. For cooling, culture plates were taken out from the home incubator (37°C) and immediately transferred to another incubator set at 31°C for 5 hours.

### Blood analyses

Human and mouse plasma ANP and BNP were measured with Human Atrial Natriuretic Peptide ELISA kit (Cusabio) and Mouse Atrial Natriuretic Peptide ELISA kit (Cusabio), Human Brain Natriuretic Peptide ELISA kit (Cusabio), and Mouse Brain Natriuretic Peptide ELISA kit (Cusabio), respectively following manufactory instructions. Glycerol was measured by enzymatic assay (Free Glycerol reagent, Sigma), NEFA and TG were measured using the NEFA C kit (Wako) and TG reagent (sigma). Glucose and ketone bodies levels were measured using a glucometer (Accucheck; Roche, Meylan, France) and blood β-ketone bodies meter (Freestyle Optium H meter, Abbot) respectively.

### Echocardiography

Echocardiography was carried out with a Vivid7 echograph (GE Healthcare) and a 14 MHz transducer (i13L, GE) on lightly anesthetized (1% isoflurane in air) mice placed on a heating pad. Left ventricular walls and cavity dimensions were obtained from parasternal short axis view at mid-ventricular level during Time Movement mode acquisition. LV mass was estimated by a spherical approximation. LV ejection fraction was measured from parasternal long axis view by delineating LV chamber area in diastole and systole. The operator was blind from mice genotype.

### ^18^F-FDG PET/CT

Positron emission tomography-computed tomography imaging with [^18^F]fluorodeoxyglucose (^18^F-FDG PET/CT) was performed as previously described ^50^. Briefly, mice were placed singly in cages, with water but without food and bedding, for 4h fasting either at 4°C or RT before transfer to the imaging lab. Then mice were injected intraperitoneally with 10 to 14.5 MBq of ^18^F-FDG (GlucoTep® Cyclopharma, S^t^-Beauzire, France, #FDGTCPRECH) and placed back into their respective cage either kept on ice or remaining on a heating pad (RT) for 1h. After anesthesia with 4% isoflurane, mice were placed in 36°C imaging chambers for a 15 min PET acquisition (NanoScan PET/CT *Mediso Ltd, Hungary*) 1h post ^18^F-FDG injection and for a 6 min CT scan imaging (720 projections, semi-circular scan method, X ray energy: 35kVp, exposure time: 450ms, voxel size: 251 x 251 x 251µm). PET acquisitions were performed in list-mode and reconstructed with a three-dimensional iterative algorithm (Tera-Tomo 3D, full detector model and low regularization; *Mediso Ltd, Hungary*) with four iterations and six subsets and a voxel size of 0.4 x 0.4 x 0.4 mm. All images were automatically corrected for radioactive decay during acquisition by the manufacturer software setting (Nucline, *Mediso Ltd, Hungary*). CT images were automatically fused to PET images and were also used for attenuation correction of PET images during their reconstruction. After acquisition, mice were placed back in clean cages with free access to food and water at RT. Processing of reconstructed images has been performed with VivoQuant software (InviCRO). 3D volumes of interest (VOIs) were drawn manually on the CT (part of left quadriceps) giving access to muscle mean ^18^F-FDG uptake (kBq.g^-1^) or, for BAT, by semi-automatic segmentation based on connected pixels threshold to calculate BAT ^18^F-FDG uptake (kBq) and metabolic volume (mm^3^).

### Adipocyte lipolysis

Fresh eWAT pads were dissected from mice and put into phosphate-buffered saline (PBS) at 37°C. Adipose tissue depots were cut into small pieces and transferred into Krebs-Ringer bicarbonate buffer (pH 7.4) containing 0.5 mM CaCl2, 238 mg/100ml HEPES, 108 mg/100 ml glucose, 3.5% BSA, and 1 mg/ml collagenase (Sigma) at 37°C for 20 min. At the end of digestion, the fat cell suspension was filtered and rinsed three-times. Isolated packed adipocytes were diluted to 1/10^th^ and incubated in Krebs-Ringer bicarbonate for basal lipolysis determination and 1µM isoproterenol (Sigma,) or 1µM ANP (Sigma) for lipolysis determination. Incubations were carried out for 90 min at 37°C and then placed on ice to stop lipolysis. Glycerol was measured by enzymatic assay (Free Glycerol reagent, Sigma,) and NEFA were measured using the NEFA C kit (Wako).

### Histology

Adipose tissues and liver were fixed with 4% paraformaldehyde in PBS, dehydrated, embedded in paraffin, and cut into 7µm sections. Sections were stained with hematoxylin and eosin using standard protocols.

### Immunofluorescence

iBAT sections (300 μm) were incubated in blocking solution (2% normal horse serum and 0.2% triton X-100 in PBS) for 4 hours at RT and then incubated with the lipid probe BODIPY 558/568 C12 (1:1000) before nuclei being stained with DAPI (Sigma). iBAT sections were also incubated with lectin-rhodamine (1:250, Vector Laboratories) overnight at 4°C and nuclei stained with DAPI. Imaging was performed using a confocal Laser Scanning microscope (LSM 780, Carl Zeiss) and image analysis was performed using Fiji software (NIH).

### G6Pase activity

Frozen tissues were homogenized using Fast Prep^®^ in 10 mM HEPES and 0.25 M sucrose, pH 7.4 (9 vol./g tissue). G6Pase activity was assayed in homogenates for 10 min at 30°C at pH 7.3 in the presence of a saturating glucose-6-phosphate concentration (20 mM) ^51^.

### Hepatic glycogen content

Liver samples were weighed and homogenized in acetate buffer (0,2M, pH 4.8). After centrifuging the samples at 12 000g for 10min, supernatant was transferred into clean tubes and divided in two aliquots. An aliquot of each homogenate was mixed with amyloglucosidase (Sigma) and incubated at 55°C for 15 minutes. The other one was mixed with water and incubated at RT for 15 minutes. Glucose content was measured as previously described below. Samples were analyzed in duplicate and the results determined as μg glycogen per mg tissue.

### Hepatic triglycerides content

Liver triglycerides were extracted using Folch extraction procedure, as previously described ^52^. This procedure consist in the addition of a chloroform/methanol 2/1 solution to 100 mg of frozen liver (1.7mL for 100mg of tissue), and then a crushing with Fast Prep^®^. The solution was centrifuged twice (2,000g; 10 min; 4°C), and 2mL of NaCl was added to the supernatant previously removed. Two phases were created, and the inferior organic phase that contains triglycerides was kept. After chloroform evaporation, triglycerides were diluted in 100µL of propanol and measured with a colorimetric kit (DiaSys, Holzheim, Germany).

### SeaHorse

For oxygen consumption measurements, differentiated adipocytes were incubated for 1 h at 37°C in unbuffered XF assay medium supplemented with 2 mM GlutaMAX, 1 mM sodium pyruvate, and 25 mM glucose. To determine mitochondrial uncoupling, oxygen consumption was measured using bio-analyzer from Seahorse Bioscience after addition of 2 µM oligomycin, which inhibited ATPase, followed by indicated concentrations of ANP or 1 µM NE. Maximal respiration was determined following 0.3 µM FCCP. 1 mM antimycin A and rotenone was added to correct for non-mitochondrial respiration ^42^.

### Real-time qPCR

Total RNA from tissue or cells was isolated using Qiagen RNeasy kit (Qiagen, GmbH Hilden, Germany) following manufacturer’s protocol. The quantity of the RNA was determined on a Nanodrop ND-1000 (Thermo Scientific, Rockford, IL, USA). Reverse-transcriptase PCR was performed using the Multiscribe Reverse Transcriptase method (Applied Biosystems, Foster City, CA). Quantitative Real-time PCR (qRT-PCR) was performed in duplicate using the ViiA 7 Real-time PCR system (Applied biosystems). All expression data were normalized by the 2^(-ΔCt)^ method using *18S* in mice and mouse cultures and *PUM1* and *GUSB* in human cultures, as internal control. Correlation with thermogenic markers gene expression was assessed using the Biomark HD system with 96 Dynamic Array IFC (Fluidigm) and TaqMan assays (Applied Biosystems) as described in ^24^. Data were normalized using the 2^-ΔCt^ method and *PUM1* as reference gene. Primer sequences are listed in **Supplementary Table 1**.

### Western blot

Proteins were extracted from tissues using Ripa buffer and protease inhibitor cocktail (Sigma-Aldrich). Tissues homogenates were centrifuged twice for 20 min at 12700 rpm and supernatants were quantified with BCA pierce kit (ThermoScientific). Equal amount of proteins were run on a 4-20% SDS-polyacrylamide gel electrophoresis (Biorad), transferred onto nitrocellulose membrane (Bio-Rad) and incubated overnight at 4°C with primary antibodies, Rabbit anti-ATLG (1:1000, CST #2138s), Rabbit anti-GAPDH (1:1000, CST, #2118), Rabbit anti-pS660 HSL (1:1000, CST, #4126), Rabbit anti-pS563 HSL (1:1000, CST, #4139s), Rabbit anti-HSL (1:1000, CST, #4107), Rabbit anti-NPRA (1:1000, Abcam, ab154266), Goat anti-NPRC (1 :1000, Sigma, SAB2501867), Rabbit anti-p-p38MAPK (1:1000, CST, #9211), Rabbit anti-P38MAPK (1:1000, CST, #9212), Rabbit anti-UCP1 (1:1000, Abcam, ab10983), Rabbit anti-G6PC^53^ (1:2000), Rabbit anti-PEPCK (1:7000, Santa cruz, #32879), Rabbit anti-Actin (1 :10000, CST, #4970). Subsequently, immuno-reactive proteins were blotted with anti-rabbit or goat horseradish peroxidase-labeled secondary antibodies for 1h at room temperature and revealed by enhanced chemiluminescence reagent (SuperSignal West Femto, Thermo Scientific), visualized using ChemiDoc MP Imaging System and data analyzed using the ImageLab 4.2 version software (Bio-Rad Laboratories, Hercules, USA).

### Statistical analyses

All statistical analyses were performed using GraphPad Prism 7.0 for Windows (GraphPad Software Inc., San Diego, CA), except for Figure 4A that was produced using the package corrplot of the R software ^10, 24^. Normal distribution and homogeneity of variance of the data were tested using Shapiro-Wilk and F tests, respectively. Student’s *t-*tests, Mann-Whitney test or one-way ANOVA were performed to determine differences between groups/treatments. Two-way ANOVA followed by Bonferonni’s post hoc tests were applied when appropriate. Univariate linear regressions were performed on parametric data. The false discovery rate for multiple testing was controlled by the Benjamini-Hochberg procedure with p_adj._ values ≤ 0.05. as threshold. All values in Figures are presented as mean ± SEM. Statistical significance was set at *P* < 0.05.

## Data availability

The data that support the findings of this study are available from the corresponding author upon reasonable request.

## Acknowledgements

This work was supported by grants from Inserm, Paul Sabatier University, Société Francophone du Diabète and European Foundation for the Study of Diabetes (C.M.), Commission of the European Communities (FP6-513946 DiOGenes and HEALTH-F2-2011-278373 DIABAT to D.L.). D.C. is supported by a Ph.D. fellowship from Inserm/Occitanie Region. We are very grateful to Caroline Nevoit (ENI CREFRE) and to Sarah Gandarillas and Candy Escassut (Animal Care Facility CREFRE) for technical assistance in PET/CT imaging. We also thank Alexandre Lucas (APC core facility), Frédéric Martins (GET-TQ core facility), and Lucie Fontaine (Histology core facility) for their technical support. We are also grateful to the study participants in clinical studies. D.L. is a member of Institut Universitaire de France. We warmly acknowledge Pr. Max Lafontan for critical reading of the manuscript.

## Author contributions

Conceptualization, D.C. and C.M.; Methodology, D.C. and C.M.; Investigation, D.C., M.C., E.N., V.B., D.L., C.P., C.L., J.VP., M.S., L.M., MA.M., Y.J., Y.SM., A.M., S.D., G.T., N.V., V.B., F.L., A.C., WHM.S., A.A.; Resources, G.M., W.VM.L., P.S., D.L.; Writing – Original Draft, D.C. and C.M.; Writing – Review & Editing, D.C., E.N., A.C., L.C., G.M., W.VM.L., P.S., D.L., and C.M.; Supervision, D.C. and C.M.; Funding Acquisition, D.L. and C.M.

## References

1. Nedergaard, J. & Cannon, B. The changed metabolic world with human brown adipose tissue: therapeutic visions. Cell metabolism 11, 268–272 (2010).

2. Kajimura, S., Spiegelman, B.M. & Seale, P. Brown and Beige Fat: Physiological Roles beyond Heat Generation. Cell metabolism 22, 546–559 (2015).

3. Cannon, B. & Nedergaard, J. Brown adipose tissue: function and physiological significance. Physiological reviews 84, 277–359 (2004).

4. Harms, M. & Seale, P. Brown and beige fat: development, function and therapeutic potential. Nature medicine 19, 1252–1263 (2013).

5. Wu, J., et al. Beige adipocytes are a distinct type of thermogenic fat cell in mouse and human. Cell 150, 366–376 (2012).

6. Heine, M., et al. Lipolysis Triggers a Systemic Insulin Response Essential for Efficient Energy Replenishment of Activated Brown Adipose Tissue in Mice. Cell metabolism 28, 644–655 e644 (2018).

7. Jiang, H., Ding, X., Cao, Y., Wang, H. & Zeng, W. Dense Intra-adipose Sympathetic Arborizations Are Essential for Cold-Induced Beiging of Mouse White Adipose Tissue. Cell metabolism 26, 686–692 e683 (2017).

8. Cao, W., et al. p38 mitogen-activated protein kinase is the central regulator of cyclic AMP-dependent transcription of the brown fat uncoupling protein 1 gene. Molecular and cellular biology 24, 3057–3067 (2004).

9. Simcox, J., et al. Global Analysis of Plasma Lipids Identifies Liver-Derived Acylcarnitines as a Fuel Source for Brown Fat Thermogenesis. Cell metabolism 26, 509–522 e506 (2017).

10. Ikeda, K., et al. UCP1-independent signaling involving SERCA2b-mediated calcium cycling regulates beige fat thermogenesis and systemic glucose homeostasis. Nature medicine 23, 1454–1465 (2017).

11. de Jong, J.M.A., et al. The beta3-adrenergic receptor is dispensable for browning of adipose tissues. American journal of physiology. Endocrinology and metabolism 312, E508–E518 (2017).

12. Bachman, E.S., et al. betaAR signaling required for diet-induced thermogenesis and obesity resistance. Science 297, 843–845 (2002).

13. Kuhn, M. Molecular Physiology of Membrane Guanylyl Cyclase Receptors. Physiological reviews 96, 751–804 (2016).

14. Moro, C., et al. Atrial natriuretic peptide contributes to physiological control of lipid mobilization in humans. FASEB journal : official publication of the Federation of American Societies for Experimental Biology 18, 908–910 (2004).

15. Sengenes, C., Berlan, M., De Glisezinski, I., Lafontan, M. & Galitzky, J. Natriuretic peptides: a new lipolytic pathway in human adipocytes. FASEB journal : official publication of the Federation of American Societies for Experimental Biology 14, 1345–1351 (2000).

16. Bordicchia, M., et al. Cardiac natriuretic peptides act via p38 MAPK to induce the brown fat thermogenic program in mouse and human adipocytes. The Journal of clinical investigation 122, 1022–1036 (2012).

17. Zhang, F., et al. An Adipose Tissue Atlas: An Image-Guided Identification of Human-like BAT and Beige Depots in Rodents. Cell metabolism 27, 252–262 e253 (2018).

18. Yuan, K., et al. Modification of atrial natriuretic peptide system in cold-induced hypertensive rats. Regulatory peptides 154, 112–120 (2009).

19. Gnad, T., et al. Adenosine activates brown adipose tissue and recruits beige adipocytes via A2A receptors. Nature 516, 395–399 (2014).

20. Mallela, J., et al. Natriuretic peptide receptor A signaling regulates stem cell recruitment and angiogenesis: a model to study linkage between inflammation and tumorigenesis. Stem Cells 31, 1321–1329 (2013).

21. Mandard, S., Muller, M. & Kersten, S. Peroxisome proliferator-activated receptor alpha target genes. Cellular and molecular life sciences : CMLS 61, 393–416 (2004).

22. de la Rosa Rodriguez, M.A. & Kersten, S. Regulation of lipid droplet-associated proteins by peroxisome proliferator-activated receptors. Biochimica et biophysica acta. Molecular and cell biology of lipids 1862, 1212–1220 (2017).

23. Puigserver, P., et al. A cold-inducible coactivator of nuclear receptors linked to adaptive thermogenesis. Cell 92, 829–839 (1998).

24. Coue, M., et al. Natriuretic peptides promote glucose uptake in a cGMP-dependent manner in human adipocytes. Scientific reports 8, 1097 (2018).

25. Kovacova, Z., et al. Adipose tissue natriuretic peptide receptor expression is related to insulin sensitivity in obesity and diabetes. Obesity (Silver Spring) 24, 820–828 (2016).

26. Ryden, M., et al. Impaired atrial natriuretic peptide-mediated lipolysis in obesity. Int J Obes (Lond) 40, 714–720 (2016).

27. Coue, M., et al. Defective Natriuretic Peptide Receptor Signaling in Skeletal Muscle Links Obesity to Type 2 Diabetes. Diabetes 64, 4033–4045 (2015).

28. Frontini, A. & Cinti, S. Distribution and development of brown adipocytes in the murine and human adipose organ. Cell metabolism 11, 253–256 (2010).

29. Ohno, H., Shinoda, K., Spiegelman, B.M. & Kajimura, S. PPARgamma agonists induce a white-to-brown fat conversion through stabilization of PRDM16 protein. Cell metabolism 15, 395–404 (2012).

30. Jankovic, A., et al. Two key temporally distinguishable molecular and cellular components of white adipose tissue browning during cold acclimation. The Journal of physiology 593, 3267–3280 (2015).

31. Seale, P., et al. Transcriptional control of brown fat determination by PRDM16. Cell metabolism 6, 38–54 (2007).

32. Sengenes, C., et al. Natriuretic peptide-dependent lipolysis in fat cells is a primate specificity. American journal of physiology. Regulatory, integrative and comparative physiology 283, R257–265 (2002).

33. Sengenes, C., et al. Involvement of a cGMP-dependent pathway in the natriuretic peptide-mediated hormone-sensitive lipase phosphorylation in human adipocytes. The Journal of biological chemistry 278, 48617–48626 (2003).

34. Samuel, V.T. & Shulman, G.I. The pathogenesis of insulin resistance: integrating signaling pathways and substrate flux. The Journal of clinical investigation 126, 12–22 (2016).

35. Perry, R.J., et al. Hepatic acetyl CoA links adipose tissue inflammation to hepatic insulin resistance and type 2 diabetes. Cell 160, 745–758 (2015).

36. van Marken Lichtenbelt, W.D., et al. Cold-activated brown adipose tissue in healthy men. The New England journal of medicine 360, 1500–1508 (2009).

37. Blondin, D.P., et al. Dietary fatty acid metabolism of brown adipose tissue in cold-acclimated men. Nature communications 8, 14146 (2017).

38. Vosselman, M.J., et al. Low brown adipose tissue activity in endurance-trained compared with lean sedentary men. Int J Obes (Lond) 39, 1696–1702 (2015).

39. Nedergaard, J., Bengtsson, T. & Cannon, B. Unexpected evidence for active brown adipose tissue in adult humans. American journal of physiology. Endocrinology and metabolism 293, E444–452 (2007).

40. Cypess, A.M., et al. Identification and importance of brown adipose tissue in adult humans. The New England journal of medicine 360, 1509–1517 (2009).

41. Virtanen, K.A., et al. Functional brown adipose tissue in healthy adults. The New England journal of medicine 360, 1518–1525 (2009).

42. Broeders, E.P., et al. The Bile Acid Chenodeoxycholic Acid Increases Human Brown Adipose Tissue Activity. Cell metabolism 22, 418–426 (2015).

43. Vosselman, M.J., et al. Systemic beta-adrenergic stimulation of thermogenesis is not accompanied by brown adipose tissue activity in humans. Diabetes 61, 3106–3113 (2012).

44. Hassi, J., Rintamaki, H., Ruskoaho, H., Leppaluoto, J. & Vuolteenaho, O. Plasma levels of endothelin-1 and atrial natriuretic peptide in men during a 2-hour stay in a cold room. Acta physiologica Scandinavica 142, 481–485 (1991).

45. Rohm, M., et al. An AMP-activated protein kinase-stabilizing peptide ameliorates adipose tissue wasting in cancer cachexia in mice. Nature medicine 22, 1120–1130 (2016).

46. Ouellet, V., et al. Brown adipose tissue oxidative metabolism contributes to energy expenditure during acute cold exposure in humans. The Journal of clinical investigation 122, 545–552 (2012).

47. Rashed, H.M., Nair, B.G. & Patel, T.B. Regulation of hepatic glycolysis and gluconeogenesis by atrial natriuretic peptide. Archives of biochemistry and biophysics 298, 640–645 (1992).

48. Larsen, T.M., et al. Diets with high or low protein content and glycemic index for weight-loss maintenance. The New England journal of medicine 363, 2102–2113 (2010).

49. Planat-Benard, V., et al. Plasticity of human adipose lineage cells toward endothelial cells: physiological and therapeutic perspectives. Circulation 109, 656–663 (2004).

50. Wang, X., Minze, L.J. & Shi, Z.Z. Functional imaging of brown fat in mice with 18F-FDG micro-PET/CT. Journal of visualized experiments : JoVE (2012).

51. Mithieux, G., Rajas, F. & Gautier-Stein, A. A novel role for glucose 6-phosphatase in the small intestine in the control of glucose homeostasis. The Journal of biological chemistry 279, 44231–44234 (2004).

52. Monteillet, L., et al. Intracellular lipids are an independent cause of liver injury and chronic kidney disease in non alcoholic fatty liver disease-like context. Molecular metabolism 16, 100–115 (2018).

53. Rajas, F., et al. Immunocytochemical localization of glucose 6-phosphatase and cytosolic phosphoenolpyruvate carboxykinase in gluconeogenic tissues reveals unsuspected metabolic zonation. Histochemistry and cell biology 127, 555–565 (2007).

